# Non-endothelial expression of Endomucin in the mouse and human choroid

**DOI:** 10.1101/2024.03.08.584133

**Authors:** Elysse Brookins, Sophia E. Serrano, George S. Yacu, Gal Finer, Benjamin R Thomson

**Affiliations:** Department of Ophthalmology and Feinberg Cardiovascular and Renal Research Inst. Northwestern University Feinberg School of Medicine, Chicago, IL, USA; Lurie Children’s Hospital Department of Nephrology and Stanley Manne Children’s Research Inst., Chicago, IL, USA

## Abstract

Endomucin (EMCN) is a 261 AA transmembrane glycoprotein that is highly expressed by venous and capillary endothelial cells where it plays a role in VEGF-mediated angiogenesis and regulation of immune cell recruitment. However, it is better known as a histological marker, where it has become widespread due to the commercial availability of high-quality antibodies that work under a wide range of conditions and in many tissues. The specificity of EMCN staining has been well-validated in retinal vessels, but while it has been used extensively as a marker in other tissues of the eye, including the choroid, the pattern of expression has not been described in detail. Here, in addition to endothelial expression in the choriocapillaris and deeper vascular layers, we characterize a population of EMCN-positive perivascular cells in the mouse choroid that did not co-localize with cells expressing other endothelial markers such as PECAM1 or PODXL. To confirm that these cells represented a new population of EMCN-expressing stromal cells, we then performed single cell RNA sequencing in choroids from adult wild-type mice. Analysis of this new dataset confirmed that, in addition to endothelial cells, *Emcn* mRNA expression was present in choroidal pericytes and a subset of fibroblasts, but not vascular smooth muscle cells. Besides *Emcn*, no known endothelial gene expression was detected in these cell populations, confirming that they did not represent endothelial-stromal doublets, a common technical artifact in single cell RNA seq datasets. Instead, choroidal *Emcn*-expressing fibroblasts exhibited high levels of chemokine and interferon signaling genes, while *Emcn*-negative fibroblasts were enriched in genes encoding extracellular matrix proteins. *Emcn* expressing fibroblasts were also detected in published datasets from mouse brain and human choroid, suggesting that stromal *Emcn* expression was not unique to the choroid and was evolutionarily conserved. Together, these findings highlight unique fibroblast and pericyte populations in the choroid and provide new context for the role of EMCN in angiogenesis and immune cell recruitment.

## Introduction

Endomucin (Emcn) is a type 1 transmembrane sialomucin that is highly expressed by venous and capillary endothelial cells in the eye and elsewhere, as well as in skin epidermis and some populations of hematopoietic cells (1, 2). In the retina, Emcn modulates angiogenesis by facilitating VEGF-mediated phosphorylation, internalization and signaling by VEGFR2 (KDR) and loss of Emcn from endothelial cells during retinal development results in delayed vascularization consistent with reduced sensitivity to VEGF (3, 4). In addition, it acts as an anti-adhesion factor, and *Emcn* overexpression suppresses TNF-α-induced neutrophil infiltration in vessels of the ciliary body and diabetes-induced leukostasis in the retina (5, 6).

Due to its near-ubiquitous expression by capillary endothelial cells and the commercial availability of high-quality antibodies, EMCN is commonly used as an endothelial marker in tissues of the eye, kidney, lung, and elsewhere (7-12). However, its widespread use as a molecular marker has not translated to research focus, and beyond its role in VEGF signaling and neutrophil adhesion, EMCN remains poorly understood (13). Its full function in the retina remains ambiguous and its patterning and function in the rest of the eye have not been described in detail. Here, though immunostaining and analysis of a new mouse choroid single cell dataset, we report that Emcn is robustly expressed by stromal cells in the mouse and human choroid, suggesting additional roles beyond regulating VEGF signaling and endothelium-mediated leukocyte extravasation. Furthermore, these data emphasize the tissue-specific nature of sialomucin expression and show that care must be taken when using EMCN as a marker of choroidal endothelial cells.

## Results

5 μm formaldehyde-fixed paraffin sections from adult, wild-type mice were used to validate EMCN expression in the choroid of the mouse eye by immunostaining and fluorescence microscopy. As previously reported, robust EMCN expression was observed in PECAM1-positive choroidal blood vessels, with highest expression observed in the fenestrated vessels of the choriocapillaris (**Figure 1 A**). Unexpectedly, however, a population of choroidal PECAM1-negative, EMCN-positive cells were also observed that lay adjacent to choriocapillaries, and between the coriocapillaris and deeper Haller and Sattler vessels. Immunostaining of en face choroid flat mounts (**Figure 1 B** and **Supplemental figure 1**) confirmed this result, and in addition to endothelial cells of the choroiocapillaris, we observed EMCN-positive cells within the PODXL-negative spaces between choriocapillaries. This was surprising, as while endothelial EMCN expression has been extensively characterized in the retina (**Figure 1 C**) and in tissues outside of the eye, we were unable to find prior reports of fibroblast or stromal expression in any tissue.

**Figure 1.**
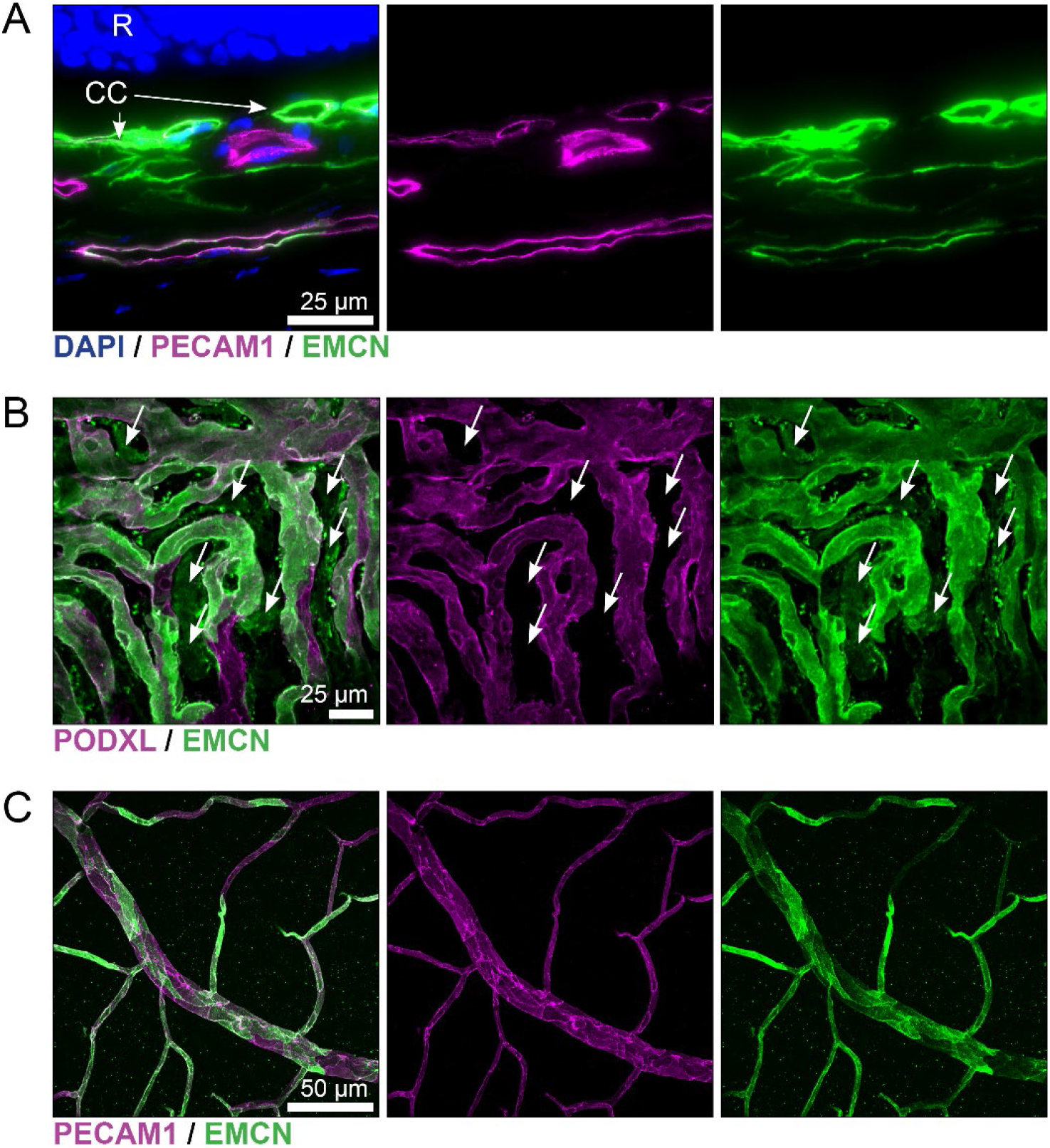
Immunostaining revealed non-endothelial endomucin (EMCN) expression in the mouse choroid, but not the retina. (A) Paraffin sections from wild-type adult mouse eyes showed that in addition to expression in the choriocapillaris (CC) and larger choroidal vessels, the mouse choroid contained extensive EMCN expression that did not colocalize with the endothelial marker PECAM1. (B) A similar result was seen in en face flat mounts, where EMCN-positive cells were seen in the area between PODXL-positive choriocapillaries (white arrows). (C) EMCN expression was confined to endothelial cells in en face retina flat mounts.

While immunostaining provided strong evidence for EMCN expression by non-endothelial cells, we turned to single cell RNA sequencing to determine the identity of this cell population and validate its existence using an independent approach. Using a variation of our previously published protocol (14), a single cell suspension was prepared from WT mouse choroids at 8 weeks of age, and single cell libraries were prepared using the 10x Genomics chromium platform. After sequencing and alignment, the Seurat package was used for analysis of the complete WT choroid dataset. We then performed quality control filtering and doublet removal before high-quality cells were projected using UMAP and clustered using a shared nearest neighbor approach. Following initial marker analysis, we manually annotated these clusters into 16 major groups (**Figure 2 A-B**). These identified categories broadly matched our expectations for choroidal cell types, although we observed some minor clusters of corneal and retinal cells that we hypothesized were the result of contamination during dissection.

**Figure 2.**
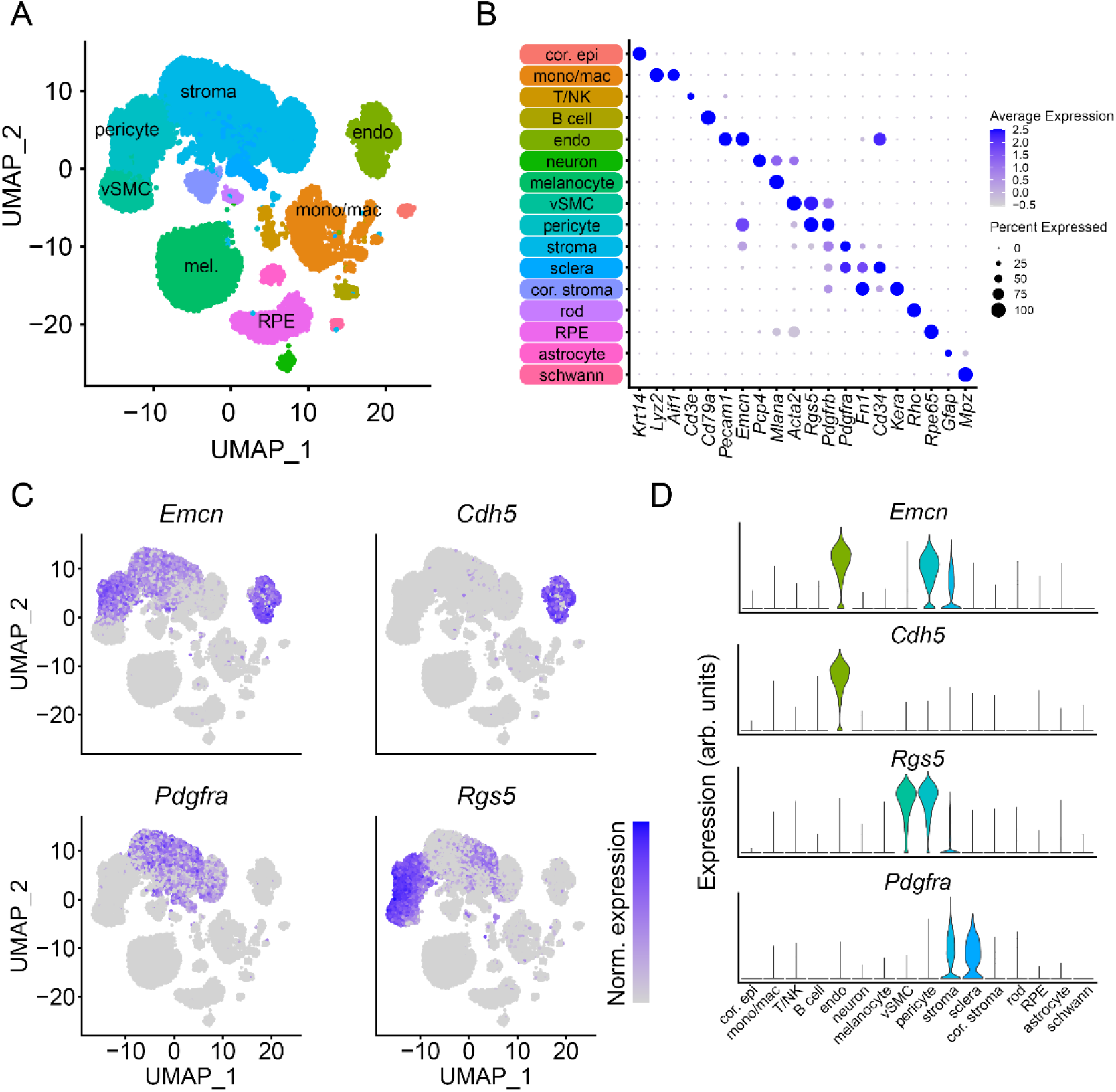
Single cell RNA sequencing confirms endomucin expression in mouse choroidal pericytes and stromal cells. (A) Single cell RNA sequencing was used to generate a dataset from wild-type mouse choroids at 8 weeks of age before cells were clustered using SEURAT and projected using UMAP. (B) Expression of known marker genes was used to annotate cell clusters and identify choroidal cell types present in the dataset. (C-D) Consistent with our immunostaining results, *Emcn* gene expression was identified in choroidal pericytes and fibroblast-like cells in addition to choroidal endothelium, but not in *Rgs5*-positive vascular smooth muscle cells.

Using our newly generated single cell dataset, we then plotted expression of *Emcn* in comparison with classical markers of endothelial cells (*Cdh5*), fibroblasts (*Pdgfra*) and vascular pericytes (*Rgs5* and *Pdgfrb-*positive; *Acta2* and *Tagln*-negative). Consistent with our immunostaining results, in addition to endothelial cells, robust *Emcn* expression was observed in *Rgs5*-positive pericytes, but not vascular smooth muscle cells, and in a subpopulation of *Pdgfra*-positive fibroblasts (**Figure 2 C-D**). As many of these cell types are closely associated with vascular endothelial cells, we then plotted expression of canonical endothelial cell markers *Kdr, Pecam1, Tek, Podxl*, and *Plvap* to exclude the possibility that the *Emcn*-positive cells we observed were pericyte/endothelium or fibroblast/endothelium doublets (**Supplemental figure 2**). While a small population of likely doublets were observed, they represented a small fraction of the *Emcn* positive cells observed in these stromal populations—confirming bona fide expression of *Emcn* in these cell types.

While our data suggested that choroidal pericytes were *Emcn*-positive, we observed distinct populations of stromal/fibroblast-like cells with differing *Emcn* expression. To aid our understanding of the role of *Emcn* in these cells, we isolated all stromal/fibroblast-like cells from our dataset and reclustered them using their own principal component space (**Figure 3 A-B**). Following re-clustering, we obtained 7 clusters in two distinct groups. Clusters 1-4 were *Emcn, Cxcl12, Cfh*-positive, while clusters 5-7 expressed *Fn1* and *Col1a1*, as well as glaucoma-related uveal gene *Myoc*. These clusters were then grouped for differential gene expression evaluation (**Figure 3 C**), followed by gene ontology analysis (**Figure 3 D**). Genes enriched in *Emcn*-positive clusters were associated with interferon signaling and neutrophil recruitment, while in contrast, *Emcn*-negative cells were enriched in genes responsible for extracellular matrix reorganization, angiogenesis and vascular development.

**Figure 3.**
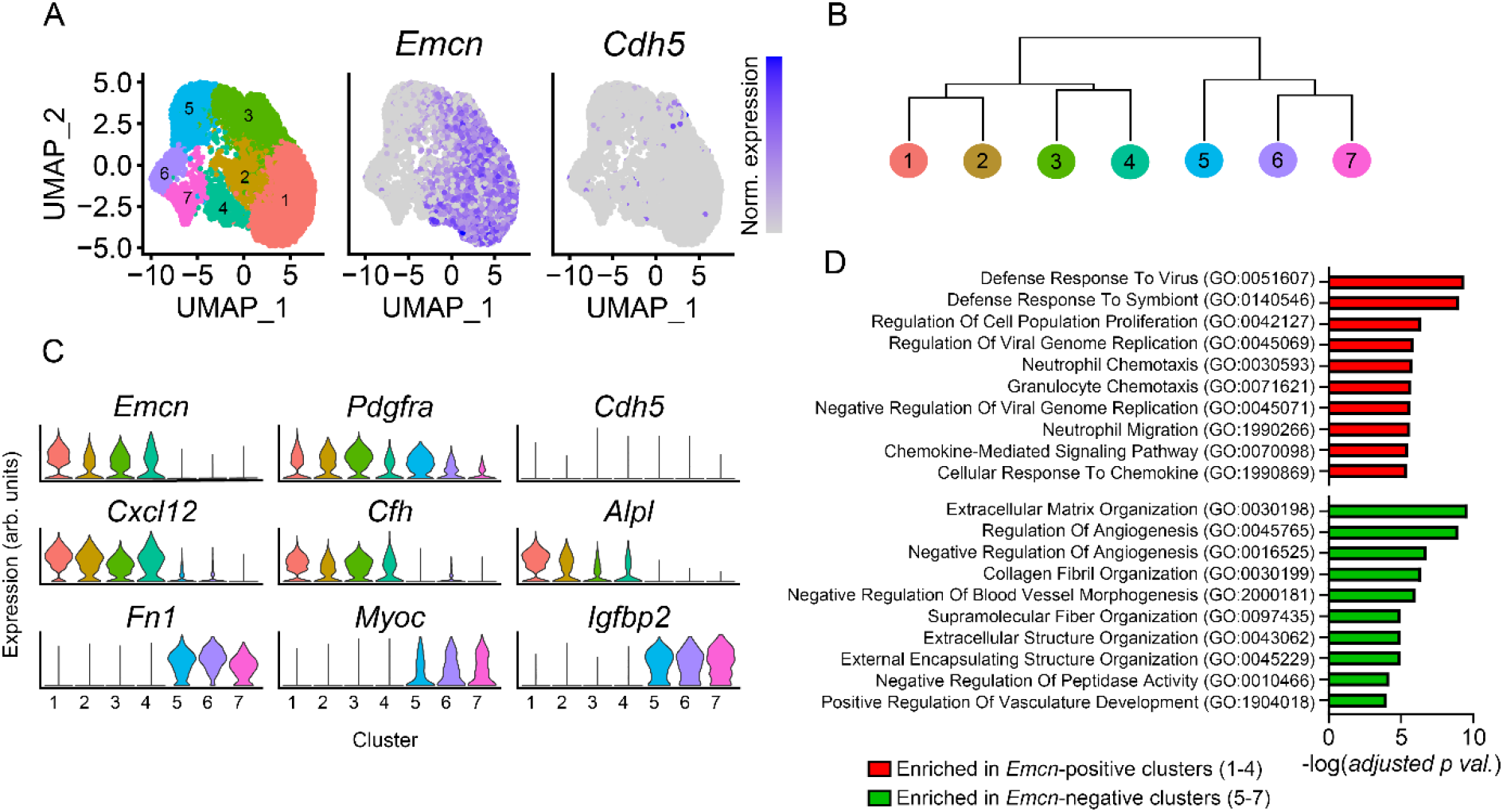
Re-clustering identifies two distinct groups of fibroblast-like choroidal stromal cells. (A-B) To determine the identify of the EMCN-positive choroidal fibroblast-like cells, we reclustered them in their own feature space. 7 clusters were identified, broadly separated into two groups. *Emcn* expression was confined to the first group, consisting of clusters 1-4. (C) This *Emcn*-positive subset was enriched in genes encoding for chemokines, including *Cxcl12*, as well as the genes encoding compliment factor H and alkaline phosphatase. In contrast, the *Emcn*-negative clusters (5-7) were enriched in genes encoding extracellular matrix proteins, the glaucoma-related gene *Myoc* and insulin-like growth factor signaling. (D) Differential gene expression between *Emcn*-expressing and *Emcn-*negative clusters was determined, and genes with a log_2_ fold change of >1 were used to perform gene ontology analysis using ENRICHR.

Seeking to confirm the presence of *Emcn*-expressing stromal cells in other tissues, we then explored a published mouse brain and lung single cell RNA seq database from the lab of Christer Betsholtz (15, 16). This data revealed two highly similar populations of brain perivascular *Pdgfra*-positive stromal cells; one highly expressing *Emcn, Cfh* and *Alpl*, and the other enriched in extracellular matrix genes such as *Fn1* (**Sup. figure 3 A**). As in our dataset, expression of *Kdr* and other endothelial cell markers was absent in these clusters, confirming that they do not represent doublets with endothelial cells. No non-endothelial expression of *Emcn* was observed in the lung (**Sup figure 4**, (15, 16)). These data suggested that *Emcn* expression in stromal cells may be widespread in tissues associated with the central nervous system. By immunostaining, while we observed robust endothelial expression, no EMCN protein was detected in non-endothelial cells of the brain (**Sup. figure 3 B**).

**Figure 4.**
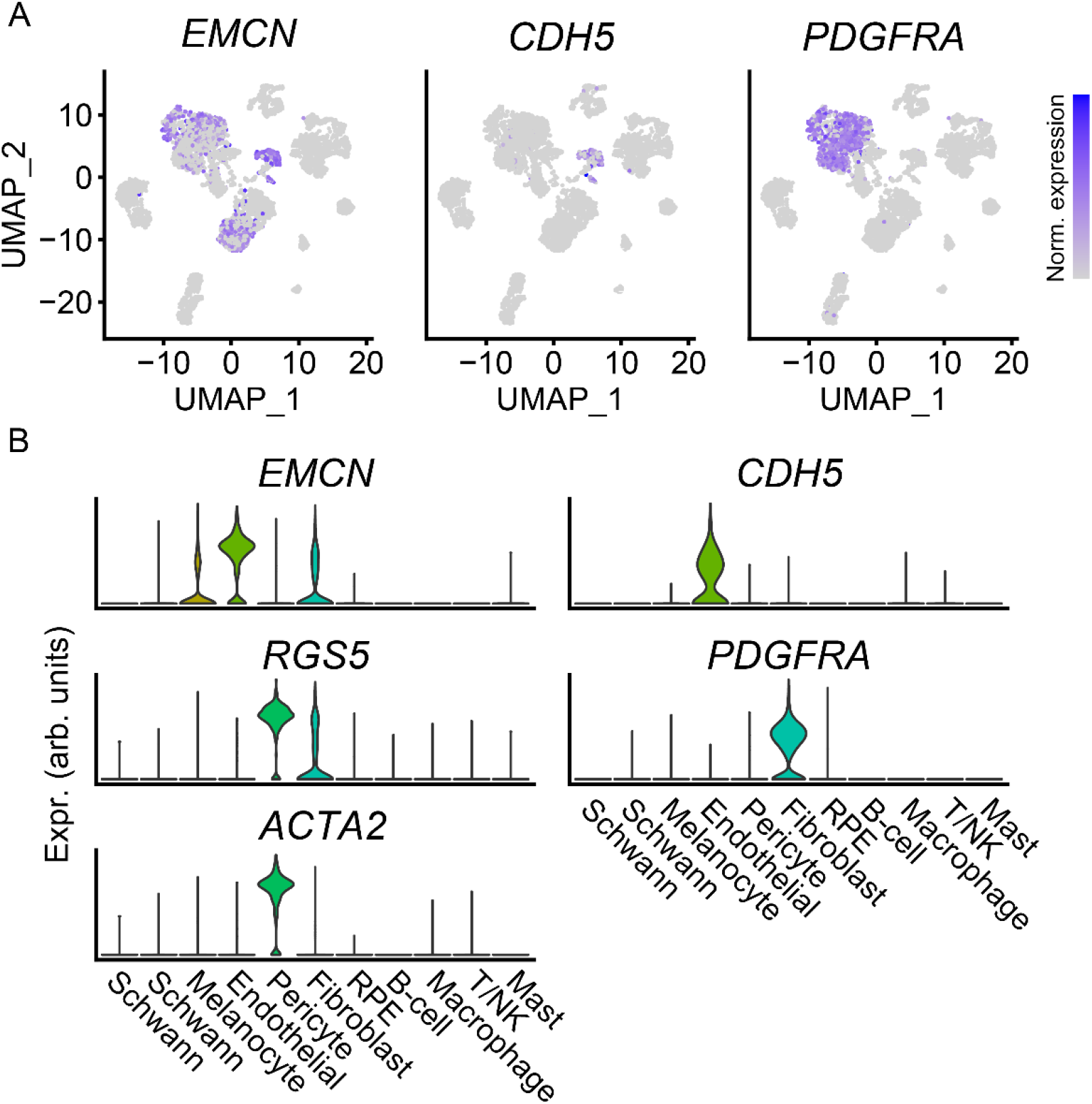
Human choroidal fibroblasts express endomucin. Our mouse findings were confirmed by re-analysis of a previously published single cell RNA sequencing dataset from the human retinal pigment epithelium and choroid (Voight et al. 2019). (A-B) In addition to *CDH5*-positive endothelial cells, *EMCN* gene expression was detected in *PDGFRA*-positive fibroblasts and some melanocytes in the human eye.

To determine if EMCN expression by choroidal pericytes and stromal cells was unique to mice, or was evolutionarily conserved, we then analyzed a published atlas of single cell RNA transcriptomics data from human choroid (17). Consistent with our findings in mouse, this dataset revealed *EMCN* expression in *PDGFRA-*positive choroidal stromal cells (**Figure 4**) although insufficient cell numbers were present in the dataset to determine if *EMCN*-positive and negative subpopulations that aligned with our mouse findings were present. No *EMCN* expression was detected in human choroidal pericytes. However, unlike our mouse dataset, cells within the human pericyte cluster expressed *ACTA2* suggesting that this cluster may include vascular smooth muscle cells in addition to true pericytes.

## Methods

### Mouse experiments

Animal studies were approved by the institutional animal care and use committee at Northwestern University and conducted in accordance with the ARVO guidelines for working with animals. Throughout the study, mice were maintained on a 12-hour light/dark cycle and provided unrestricted access to standard mouse diet and water.

### Immunostaining

Eyes were fixed (4% formaldehyde in phosphate buffered saline pH 7.5) before embedding in paraffin, sectioning and rehydrating using standard methods or processing for whole mount staining. Choroid whole mounts were bleached (3% H_2_O_2_ in phosphate buffered saline, pH 7.5, 4h at 55°) as previously described (18) before staining. After fixation, sectioning and bleaching (as appropriate), all slides and whole mounts were blocked and permeabilized (5% donkey serum, 5% bovine serum albumin, 0.5% triton X-100 in tris buffered saline pH 7.5) before incubation overnight in primary antibodies diluted in additional blocking buffer. Antibodies used: Rat anti Endomucin V.7C7 (AB106100, Abcam, Waltham MA, USA); goat anti PECAM1 (AF3628, R&D Systems, Minneapolis, MN USA); goat anti PODXL (AF1658, R&D systems). After incubation in primary antibodies, samples were washed, incubated in fluorescently labeled secondary antibodies (Life Technologies, Carlsbad CA, USA), washed a second time, and mounted for imaging. Images were captured using a Nikon W1 or Ti2 microscope and processed using ImageJ Fiji software (19).

### Single cell RNA seq

Two pooled samples, each comprised of choroids from 1 male and 1 female WT mouse were prepared using a modified version of our previously published protocol for preparation of samples from the anterior segment (14). Briefly, mice were euthanized before eyes were enucleated and choroids dissected in ice-cold HBSS (Gibco-Life Technologies, Carlsbad CA) under a dissecting microscope. After dissection, pooled choroids were minced and immediately transferred to ice cold DMEM (Gibco) containing 10% fetal bovine serum. Samples were then digested for 2h in 1mg/ml Collagenase A (Roche, Basel, Switzerland) solution containing 1 μM of Y27362 (Roche), washed and digested for an additional 30 minutes in 0.25% trypsin (Gibco) containing 100 units/ml DNAse 1 (Roche). Samples were then washed a second time, passed through a 40 μm filter and submitted to the NU Seq core at Northwestern University for single cell library preparation using the 10x chromium platform. Libraries were then sequenced using a 50 bp single end protocol on an Illumina NovaSeq platform.

### Bioinformatics analysis

Sequencing results were aligned to the mm10 reference assembly using 10x Genomics CellRanger and analyzed using Seurat 4.3.0 in R (20). The SoupX package was used to remove ambient RNA contamination (21), and the data was filtered to exclude features detected in <2 droplets per sample, droplets containing <500 or >5000 unique features, or droplets with >10% reads mapping to mitochondrial genes. Predicted doublets were then removed using the scDblFinder package (22) before sample integration and analysis in Seurat. Samples were integrated using an anchor-based CCA pipeline and the 5000 most highly variable features in the dataset. Genes which were detected in only a single sample were excluded from the anchor set. PCA was performed on the integrated dataset in 50 dimensions, and UMAP was used to visualize PCA-reduced data (min.dist = 1.5, n.neighbors = 800). Clustering was then performed in Seurat using a Louvain algorithm at a resolution of 0.6, resulting in 28 unique clusters. Clusters were manually annotated using differentially expressed genes to identify the 16 broad cell categories described in the manuscript.

Reclustering of stromal cells was performed using a new feature space of 2000 highly variable genes. 10 principal components were calculated within this feature space and used for UMAP (min.dist = 0.5, n.neighbors = 800), and clustering at a resolution of 0.4. Differentially expressed genes were then analyzed using the FindMarkers() function in Seurat and the MAST package (23). For gene ontology analysis, differentially expressed genes with a log_2_ fold change of > 1 were used as input for ENRICHR (https://maayanlab.cloud/Enrichr/, (24)). The top 10 GO terms were reported in the manuscript.

Code used for analysis and to generate figures is available via github (URL to be provided upon publication).

## Discussion

Despite our still limited understanding of its functional role, Emcn has arisen as a useful marker of capillary and venous endothelial cells due to its robust cell-surface expression and the commercial availability of high-quality antibodies. Endothelial-specific expression has been widely demonstrated in tissues of the kidney, liver, heart, lung and eye, as well as various tumors (25). Non endothelial expression has been reported in skin (1), and in hematopoietic stem cells (26). In the mouse choroid, we and others have reported high EMCN expression that was attributed to endothelial cells (8-10). However, while EMCN expression is indeed high in choriocapillaries and other endothelial cells, at least some expression in these tissues originates in pericytes and fibroblasts. In light of this finding, we believe that other groups have also observed non-endothelial choroidal EMCN expression, although it has not always been recognized. For example, immunostaining of mouse choroids by Saint-Geniez et al. demonstrated that while PECAM1 staining was clearly confined to the choriocapillaris and deeper vascular layers, EMCN expression was ubiquitous throughout the choroid (9). We believe that these previous findings are consistent with our results, although no double labeling was reported.

EMCN is well-validated as a specific marker of endothelial cells in the retina. As we did not detect non-endothelial EMCN protein expression in that tissue, we were surprised to find *Emcn* mRNA expression in a published dataset of *Pdgfra*-positive mouse brain fibroblasts, suggesting that fibroblast *Emcn* expression is present in tissues of the central nervous system and is not unique to the choroid. However, despite high mRNA observed by single cell transcriptomics, we only detected EMCN protein in brain endothelial cells and did not observe expression in other cell types. While we cannot rule out the possibility that EMCN protein is not present in these cells, our experiments were performed using the monoclonal antibody V.7C7, which recognizes a glycosylated form of EMCN and is sensitive to sialidase treatment (2). It is possible that non-endothelial EMCN expressed in tissues of the CNS are differentially glycosylated compared to those in the choroid, resulting in loss of antigenicity using the V.7C7 monoclonal. A similar finding has been described in endothelial cells of high endothelial venules, where anti-mouse monoclonal antibodies (including V.7C7) failed to detect EMCN, while robust expression was detected using polyclonal antibodies that were agnostic to glycosylation pattern (1, 2). Similarly, we did not observe perivascular EMCN expression in the retina and were unable to explore *Emcn* mRNA expression in retinal pericytes. Our single cell dataset contains only choroidal tissue and previously published datasets we analyzed contained either insufficient numbers of pericytes or high numbers of endothelial-pericyte doublets (characterized by co-detection of multiple pericyte and endothelial markers in the same droplet). Additional studies using larger datasets of retinal pericytes or antibodies that are not affected by glycosylation are required to conclusively validate expression (or lack thereof) in this tissue.

Given the near-ubiquitous *Emcn* mRNA expression we observed in the mouse choroidal pericyte cluster, we were surprised that *EMCN* was absent from human pericytes within a previously published choroidal cell atlas, despite expression in choroidal fibroblasts that was similar to our observation in mouse (17). This finding was confirmed by exploring other published human datasets accessible via *Spectacle* (27), including data from Voigt et al 2022 and Collin et al, 2023 (28, 29). Consistent with findings from our initial analysis, limited *EMCN* expression was detected in pericytes within these datasets, while fibroblast expression appeared widespread. Pericytes play a key role in regulating vascular health and function and help to mediate immune cell diapedesis (30), processes that are critical to pathogenesis of eye diseases including macular degeneration and diabetic retinopathy. If anti-adhesion molecules such as EMCN are differentially expressed between species, it could result in disease-relevant differences in immune cell response that deserve additional study. However, unlike our mouse data, the datasets of human choroidal tissue we explored reported *ACTA2* expression (encoding α smooth muscle actin) within the annotated pericyte clusters. In mouse choroid, α smooth muscle actin is confined to vascular smooth muscle and is absent from pericytes (31). A similar pattern has been reported in human tissue sections, where smooth muscle actin-positive cells are associated with larger vessels, but absent from the choriocapillaris or small vessels—consistent with the typical localization of vascular smooth muscle cells (32). It is therefore possible that annotated pericyte clusters within publicly available databases are misidentified or contain a mixture of pericytes and vascular smooth muscle cells. These closely related cell types exist in a spectrum of phenotypes and are notoriously difficult to differentiate (33). If so, this finding would be consistent with our mouse data, which demonstrates high levels of *Emcn* expression within pericytes but none in *Acta2*-positive vascular smooth muscle cells.

Together, our data demonstrate that while EMCN is a valuable marker of venous and capillary endothelium in the retina and many other tissues, it is also expressed by fibroblasts in the choroid and brain, and in mouse choroidal pericytes. Care must be taken to validate marker antibodies in each new tissue and as seen here, even within subsections of each tissue as protein localization can change and dramatically impact interpretation of results. Intriguingly, many of the key studies elucidating the role of Emcn in the eye have used whole-tissue approaches, including siRNA treatment, viral vector overexpression, and whole-body knockouts (4, 5, 34). Although the role of sialomucins in stromal tissues remains poorly understood (35), our data raises the possibility that pericyte/fibroblast EMCN may contribute to the phenotypes observed in these models, and highlights the value of future studies using tissue specific approaches.

## Acknowledgements

This work was supported by NIH R01 EY032609 and by the donors of the Brightfocus Foundation through grant M2021018N (To BRT). In addition, SS was supported by the Northwestern University Feinberg School of Medicine SciHigh program and NIH P30 DK114857 awarded to the Section of Nephrology and Hypertension. Imaging of en face choroid and retina tissue was performed at the Center for Advanced Microscopy of the Feinberg School of Medicine supported by NCI CCSG P30 CA60553. In addition, authors are indebted to Sol Misener and Kyron McAllister for technical assistance.

---

**Supplemental figure 1.**
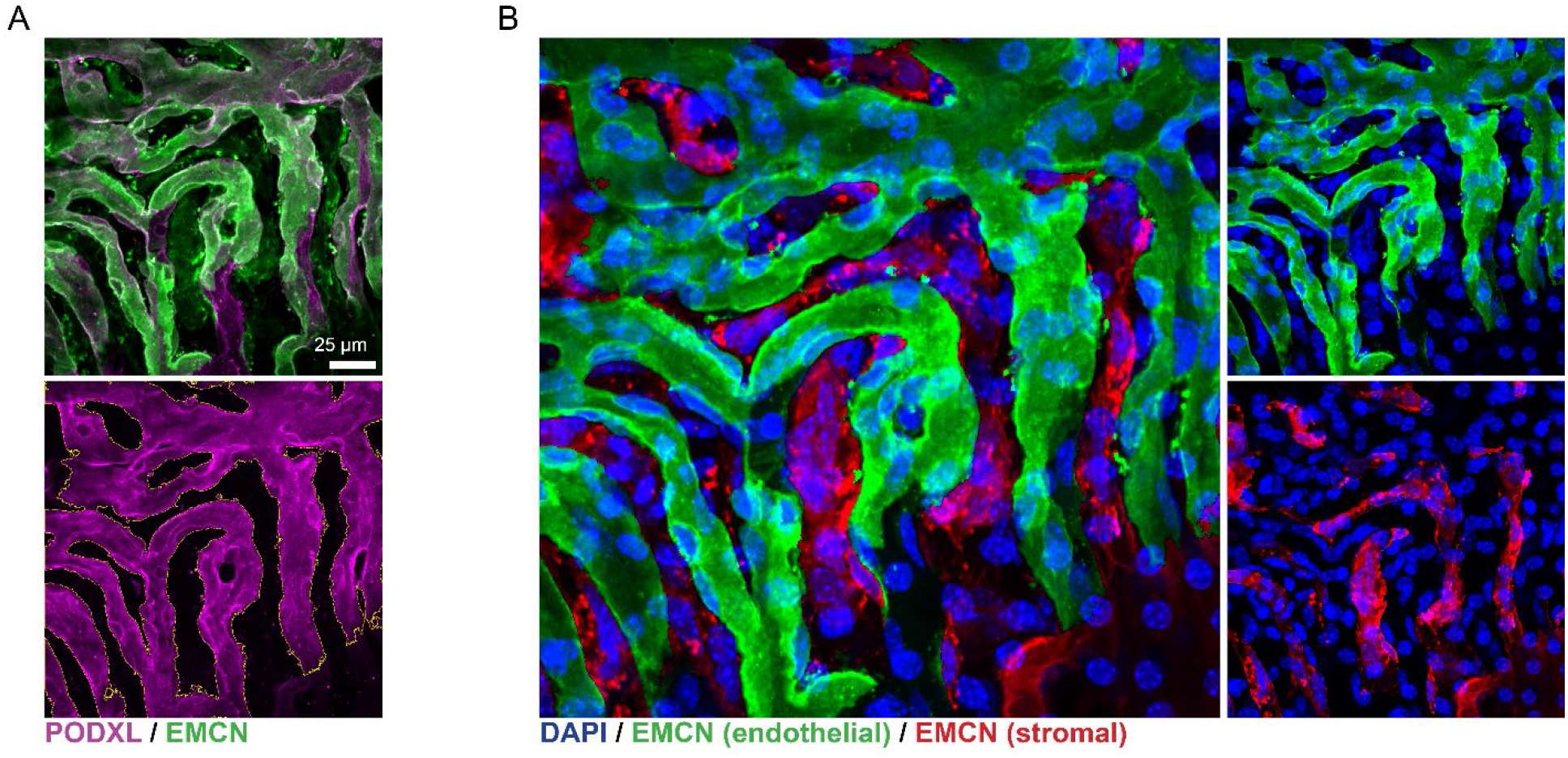
En face choriocapillaris whole mounts demonstrate non-vascular EMCN expression. (A) To clarify the expression of stromal EMCN in close proximity to high-expressing endothelial cells, PODXL staining from the whole mount image shown in figure 1 B was used to create a mask identifying endothelial cells using ImageJ Fiji. (B) The pattern of PODXL expression was then used to segment and pseudo color the EMCN staining channel of the image. In the pseudo-colored image, green represents endothelial EMCN expression, while red represents stromal expression that did not colocalize with PODXL. Note the presence of numerous DAPI-positive nuclei associated with non-endothelial EMCN expressing regions.

**Supplemental figure 2.**
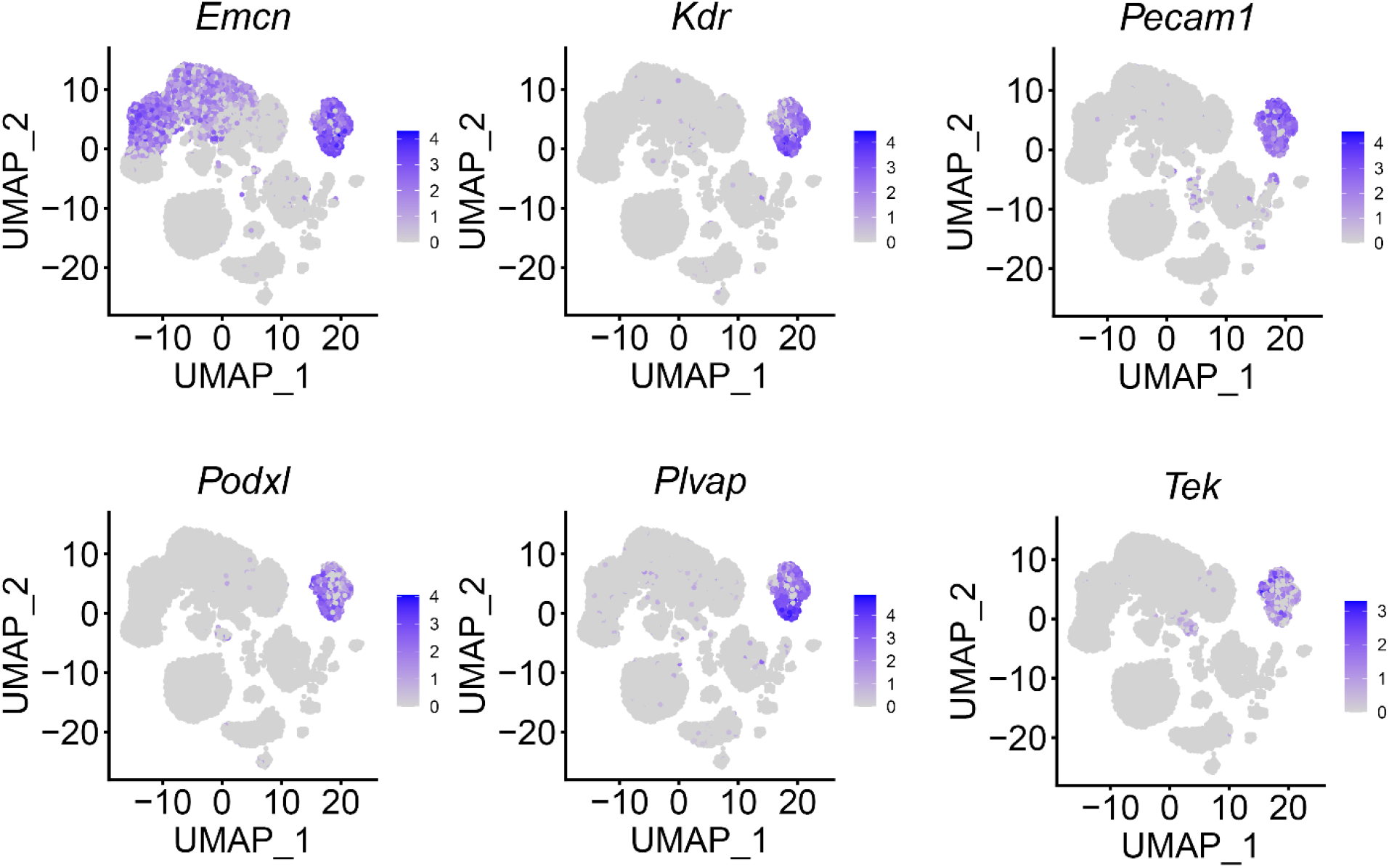
Endomucin-positive stromal cells identified by single cell RNA sequencing are not endothelial doublets. To verify that the *Emcn-*positive cell population we observed by RNA sequencing was not an artifact of endothelial-pericyte or endothelial-fibroblast doublets, we explored expression of additional endothelial markers in our dataset. Minimal expression of classical endothelial markers was observed, confirming that our detection of *Emcn* was not artifactual.

**Supplemental figure 3.**
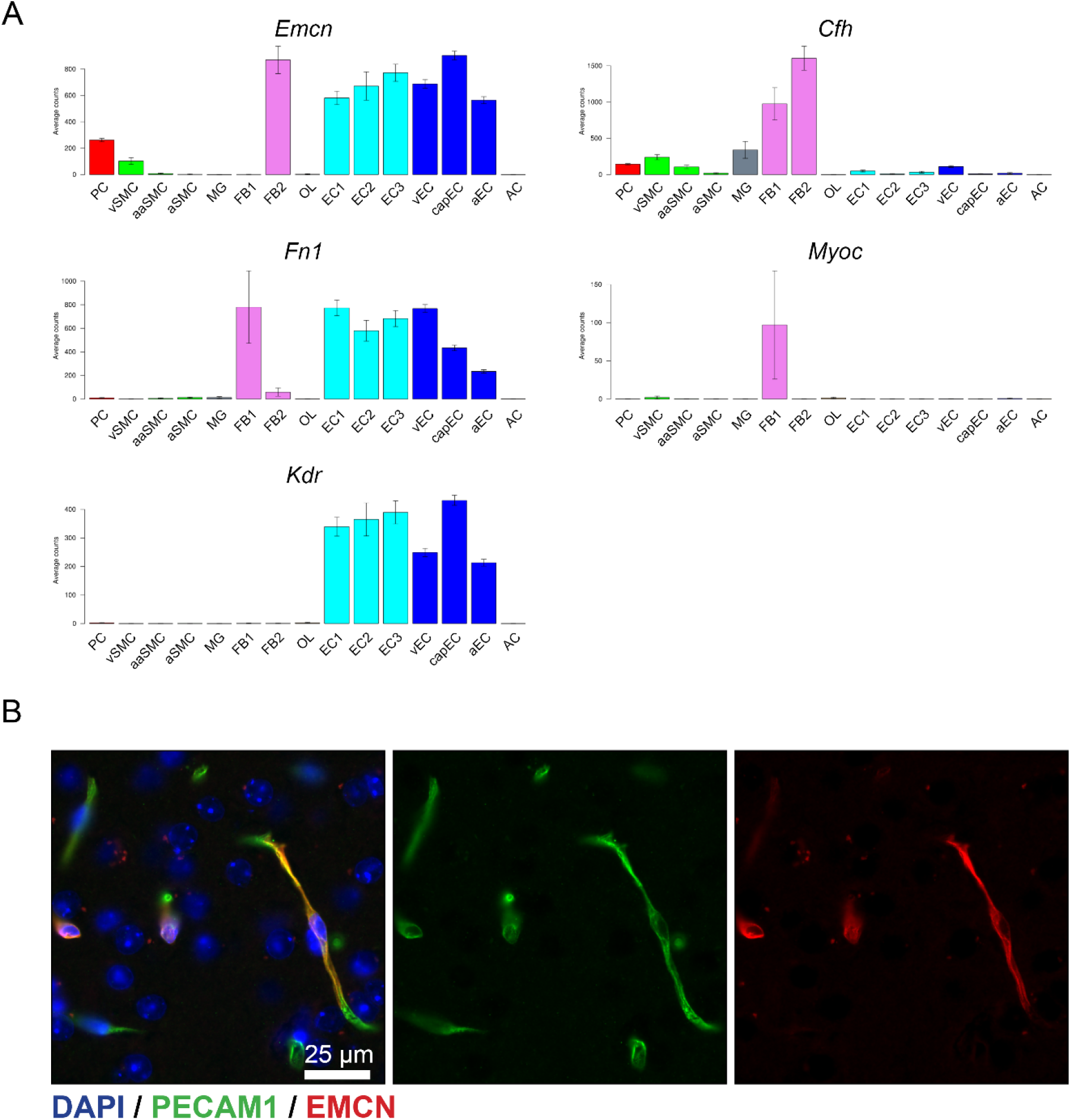
*Emcn* mRNA is expressed by *Pdgfra*-positive cells in the mouse brain but was not detected by immunostaining. (A) A previously published database of mouse brain perivascular cells (Vanlandewijck, M., He, L. et al. 2018 and He, L., Vanlandewijck, M. et al. 2018) was queried for the presence of *Emcn*-expressing pericytes and fibroblasts. This analysis revealed populations of fibroblasts that were analogous to those seen in the choroid, one *Emcn*-positive population expressing *Cfh*, and a second *Emcn*-negative population expressing extracellular matrix molecules and *Myoc*. (B). Immunofluorescence imaging of mouse brain sections did not reveal EMCN expression in non-endothelial cells.

**Supplemental figure 4.**
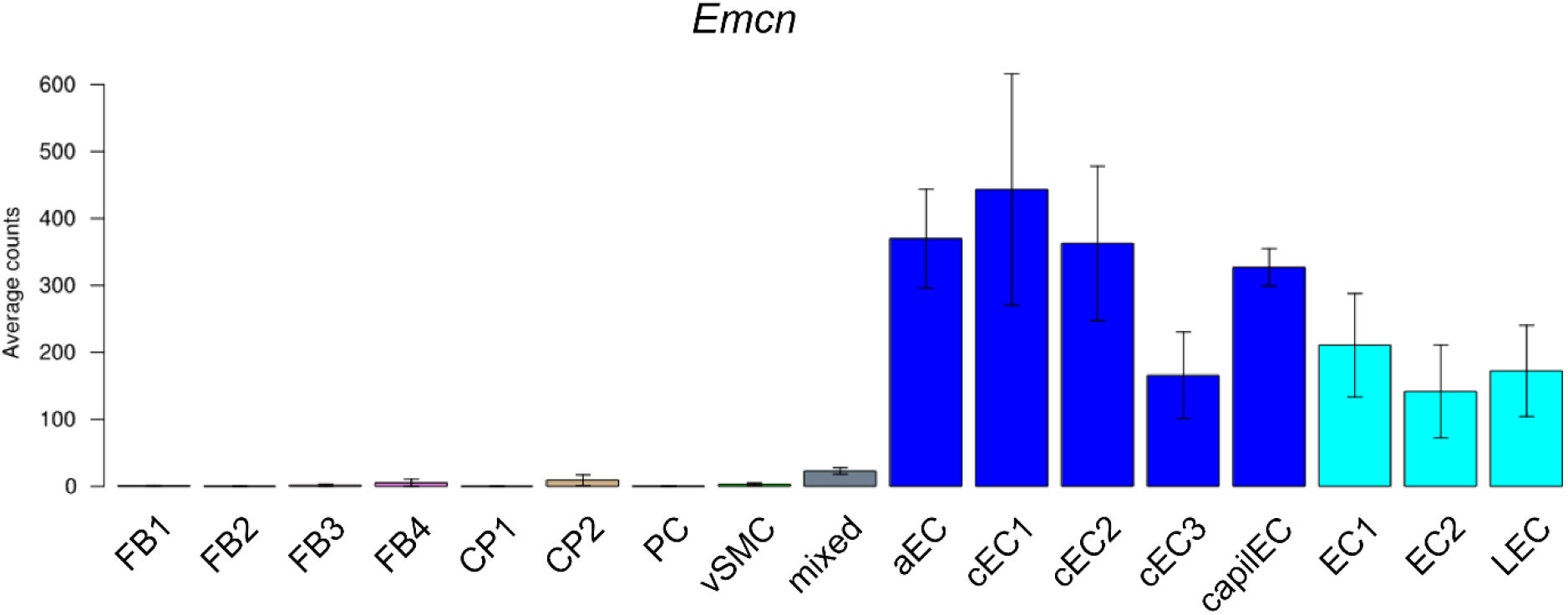
No *Emcn* expression was detected in mouse lung pericytes or fibroblasts. Analysis of a published mouse lung single cell dataset (Vanlandewijck, M., He, L. et al. 2018 and He, L., Vanlandewijck, M. et al. 2018) revealed that *Emcn* expression was confined to endothelial cells.

## References

1. Samulowitz U, Kuhn A, Brachtendorf G, Nawroth R, Braun A, Bankfalvi A, Böcker W, and Vestweber D. Human endomucin: Distribution pattern, expression on high endothelial venules, and decoration with the meca-79 epitope. Am J Pathol. 2002;160(5):1669–1669. DOI: 10.1016/s0002-9440(10)61114-5

2. Morgan SM, Samulowitz U, Darley L, Simmons DL, and Vestweber D. Biochemical characterization and molecular cloning of a novel endothelial-specific sialomucin. Blood. 1999;93(1):165–165. DOI: 10.1182/blood.V93.1.165

3. Leblanc ME, Saez-Torres KL, Cano I, Hu Z, Saint-Geniez M, Ng YS, and D’amore PA. Glycocalyx regulation of vascular endothelial growth factor receptor 2 activity. FASEB J. 2019;33(8):9362–9362. DOI: 10.1096/fj.201900011R

4. Park-Windhol C, Ng YS, Yang J, Primo V, Saint-Geniez M, and D’amore PA. Endomucin inhibits vegf-induced endothelial cell migration, growth, and morphogenesis by modulating vegfr2 signaling. Sci Rep. 2017;7(1):17138. DOI: 10.1038/s41598-017-16852-x

5. Zahr A, Alcaide P, Yang J, Jones A, Gregory M, Dela Paz NG, Patel-Hett S, Nevers T, Koirala A, Luscinskas FW, et al. Endomucin prevents leukocyte–endothelial cell adhesion and has a critical role under resting and inflammatory conditions. Nature Communications. 2016;7(10363. DOI: 10.1038/ncomms10363

6. Niu T, Zhao M, Jiang Y, Xing X, Shi X, Cheng L, Jin H, and Liu K. Endomucin restores depleted endothelial glycocalyx in the retinas of streptozotocin-induced diabetic rats. The FASEB Journal. 2019;33(12):13346–13346. DOI: 10.1096/.201901161R

7. Corada M, Orsenigo F, Morini MF, Pitulescu ME, Bhat G, Nyqvist D, Breviario F, Conti V, Briot A, Iruela-Arispe ML, et al. Sox17 is indispensable for acquisition and maintenance of arterial identity. Nature Communications. 2013;4(1):2609. DOI: 10.1038/ncomms3609

8. Liu P, Lavine JA, Fawzi A, Quaggin SE, and Thomson BR. Angiopoietin-1 is required for vortex vein and choriocapillaris development in mice. Arterioscler Thromb Vasc Biol. 2022;42(11):1413–1413. DOI: doi:10.1161/ATVBAHA.122.318151

9. Saint-Geniez M, Maldonado AE, and D’amore PA. Vegf expression and receptor activation in the choroid during development and in the adult. Invest Ophthalmol Vis Sci. 2006;47(7):3135–3135. DOI: 10.1167/iovs.05-1229

10. Goto S, Onishi A, Misaki K, Yonemura S, Sugita S, Ito H, Ohigashi Y, Ema M, Sakaguchi H, Nishida K, et al. Neural retina-specific aldh1a1 controls dorsal choroidal vascular development via sox9 expression in renal pigment epithelial cells. eLife. 2018;7(e32358. DOI: 10.7554/eLife.32358

11. Dimke H, Sparks MA, Thomson BR, Frische S, Coffman TM, and Quaggin SE. Tubulovascular cross-talk by vascular endothelial growth factor a maintains peritubular microvasculature in kidney. Journal of the American Society of Nephrology. 2014. DOI: 10.1681/asn.2014010060

12. Lange AW, Haitchi HM, Lecras TD, Sridharan A, Xu Y, Wert SE, James J, Udell N, Thurner PJ, and Whitsett JA. Sox17 is required for normal pulmonary vascular morphogenesis. Dev Biol. 2014;387(1):109–109. DOI: 10.1016/j.ydbio.2013.11.018

13. Zhang G, Yang X, and Gao R. Research progress on the structure and function of endomucin. Animal Model Exp Med. 2020;3(4):325–325. DOI: 10.1002/ame2.12142

14. Thomson BR, and Quaggin SE. Preparation of a single cell suspension from the murine iridocorneal angle. Bio-protocol. 2022;12(10):e4426. DOI: 10.21769/BioProtoc.4426

15. Vanlandewijck M, He L, Mäe MA, Andrae J, Ando K, Del Gaudio F, Nahar K, Lebouvier T, Laviña B, Gouveia L, et al. A molecular atlas of cell types and zonation in the brain vasculature. Nature. 2018;554(7693):475–475. DOI: 10.1038/nature25739

16. He L, Vanlandewijck M, Mäe MA, Andrae J, Ando K, Del Gaudio F, Nahar K, Lebouvier T, Laviña B, Gouveia L, et al. Single-cell rna sequencing of mouse brain and lung vascular and vessel-associated cell types. Sci Data. 2018;5(180160. DOI: 10.1038/sdata.2018.160

17. Voigt AP, Mulfaul K, Mullin NK, Flamme-Wiese MJ, Giacalone JC, Stone EM, Tucker BA, Scheetz TE, and Mullins RF. Single-cell transcriptomics of the human retinal pigment epithelium and choroid in health and macular degeneration. Proceedings of the National Academy of Sciences. 2019;116(48):24100. DOI: 10.1073/pnas.1914143116

18. Bhutto IA, Mcleod DS, Thomson BR, Lutty GA, and Edwards MM. Visualization of choroidal vasculature in pigmented mouse eyes from experimental models of amd. Exp Eye Res. 2024;238(109741. DOI: 10.1016/j.exer.2023.109741

19. Schindelin J, Arganda-Carreras I, Frise E, Kaynig V, Longair M, Pietzsch T, Preibisch S, Rueden C, Saalfeld S, Schmid B, et al. Fiji: An open-source platform for biological-image analysis. Nat Meth. 2012;9(7):676–676. DOI: http://www.nature.com/nmeth/journal/v9/n7/abs/nmeth.2019.html#supplementary-informaon

20. Kaplan N, Wang J, Wray B, Patel P, Yang W, Peng H, and Lavker RM. Single-cell rna transcriptome helps define the limbal/corneal epithelial stem/early transit amplifying cells and how autophagy affects this population. Invest Ophthalmol Vis Sci. 2019;60(10):3570–3570. DOI: 10.1167/iovs.19-27656

21. Young MD, and Behjati S. Soupx removes ambient rna contamination from droplet-based single-cell rna sequencing data. Gigascience. 2020;9(12). DOI: 10.1093/gigascience/giaa151

22. Germain P-L, Sonrel A, and Robinson MD. Pipecomp, a general framework for the evaluation of computational pipelines, reveals performant single cell rna-seq preprocessing tools. Genome Biol. 2020;21(1):227. DOI: 10.1186/s13059-020-02136-7

23. Finak G, Mcdavid A, Yajima M, Deng J, Gersuk V, Shalek AK, Slichter CK, Miller HW, Mcelrath MJ, Prlic M, et al. Mast: A flexible statistical framework for assessing transcriptional changes and characterizing heterogeneity in single-cell rna sequencing data. Genome Biol. 2015;16(1):278. DOI: 10.1186/s13059-015-0844-5

24. Chen EY, Tan CM, Kou Y, Duan Q, Wang Z, Meirelles GV, Clark NR, and Ma’ayan A. Enrichr: Interactive and collaborative html5 gene list enrichment analysis tool. BMC Bioinformatics. 2013;14(128. DOI: 10.1186/1471-2105-14-128

25. Liu C, Shao Z-M, Zhang L, Beaty P, Sartippour M, Lane T, Livingston E, and Nguyen M. Human endomucin is an endothelial marker. Biochem Biophys Res Commun. 2001;288(1):129–129. DOI: 10.1006/bbrc.2001.5737

26. Reckzeh K, Kizilkaya H, Helbo AS, Alrich ME, Deslauriers AG, Grover A, Rapin N, Asmar F, Grønbæk K, Porse B, et al. Human adult hscs can be discriminated from lineage-committed hpcs by the expression of endomucin. Blood Adv. 2018;2(13):1628–1628. DOI: 10.1182/bloodadvances.2018015743

27. Voigt AP, Whitmore SS, Lessing ND, Deluca AP, Tucker BA, Stone EM, Mullins RF, and Scheetz TE. Spectacle: An interactive resource for ocular single-cell rna sequencing data analysis. Exp Eye Res. 2020;200(108204. DOI: 10.1016/j.exer.2020.108204

28. Voigt AP, Mullin NK, Mulfaul K, Lozano LP, Wiley LA, Flamme-Wiese MJ, Boese EA, Han IC, Scheetz TE, Stone EM, et al. Choroidal endothelial and macrophage gene expression in atrophic and neovascular macular degeneration. Hum Mol Genet. 2022;31(14):2406–2406. DOI: 10.1093/hmg/ddac043

29. Collin J, Hasoon MSR, Zer D, Hammadi S, Dorgau B, Clarke L, Steel D, Hussain R, Coxhead J, Lisgo S, et al. Single-cell rna sequencing reveals transcriptional changes of human choroidal and retinal pigment epithelium cells during fetal development, in healthy adult and intermediate age-related macular degeneration. Hum Mol Genet. 2023;32(10):1698–1698. DOI: 10.1093/hmg/ddad007

30. Joulia R, Guerrero-Fonseca IM, Girbl T, Coates JA, Stein M, Vázquez-Martínez L, Lynam E, Whiteford J, Schnoor M, Voehringer D, et al. Neutrophil breaching of the blood vessel pericyte layer during diapedesis requires mast cell-derived il-17a. Nat Commun. 2022;13(1):7029. DOI: 10.1038/s41467-022-34695-7

31. Kim SJ, Kim SA, Choi YA, Park DY, and Lee J. Alpha-smooth muscle actin-positive perivascular cells in diabetic retina and choroid. Int J Mol Sci. 2020;21(6). DOI: 10.3390/ijms21062158

32. Chan-Ling T, Koina ME, Mccolm JR, Dahlstrom JE, Bean E, Adamson S, Yun S, and Baxter L. Role of cd44+ stem cells in mural cell formation in the human choroid: Evidence of vascular instability due to limited pericyte ensheathment. Invest Ophthalmol Vis Sci. 2011;52(1):399–399. DOI: 10.1167/iovs.10-5403

33. Gerhardt H, and Betsholtz C. Endothelial-pericyte interactions in angiogenesis. Cell Tissue Res. 2003;314(1):15–15. DOI: 10.1007/s00441-003-0745-x

34. Hu Z, Cano I, Lennikov A, Wild M, Gupta U, Ng YSE, and D’amore PA. Endomucin deletion leads to disorganized choroidal capillary fenestration and reduces laser-induced choroidal neovascularization in mice. Invest Ophthalmol Vis Sci. 2023;64(8):3005-. DOI:

35. Hughes MR, Canals Hernaez D, Cait J, Refaeli I, Lo BC, Roskelley CD, and Mcnagny KM. A sticky wicket: Defining molecular functions for cd34 in hematopoietic cells. Exp Hematol. 2020;86(1–14. DOI: 10.1016/j.exphem.2020.05.004

